# Cross-Species Analysis Reveals No Universal Programmed Aging Mechanism: Insights from Single-Cell Transcriptomics in Zebrafish, Fruit Fly, and Nematode

**DOI:** 10.1101/2024.10.28.620557

**Authors:** Yunhui Niu, Dongzhi Wu, Sen Zhang, Hong Zheng, Xing Wu, Jiansong Chen, Yunze Zhang, Tao Zhang, Wenhui He, Li Chen

**Affiliations:** Institute of Orthopedics,Fuzhou Second General Hospital, Fuzhou, China; Institute of Biological Sciences, Faculty of Science, University Malaya, Kuala Lumpur, Malaysia; School of Biological and Behavioural Sciences, Queen Mary University of London, London, United Kingdom; Department of Orthopedics, Fuzhou Second General Hospital, Fuzhou, China; Department of Neurosurgery, Fuzhou Second General Hospital, Fuzhou, China; Faculty of Biology Medicine and Health, University of Manchester, Manchester, United Kingdom

## Abstract

The question of whether aging follows a universal programmed process has been a topic of debate for a long time. Previous arguments, either supporting or refuting programmed aging, were mainly based on different evolutionary biology theories. In this study, we analysed single-cell RNA sequencing data from zebrafish, fruit fly, and nematode at various stages of development to explore gene co-expression modules across these species. We successfully identified a co-expression module related to ribosomal protein genes that is shared across the early development stages in multiple tissues of all three species. However, we did not find any cross-species shared gene co-expression modules related to aging. Further analysis of gene regulatory networks (GRNs) demonstrated that although certain aging-related genes are conserved, their regulatory mechanisms vary significantly between species. These findings suggest that aging is not governed by a conserved universal program but rather by species-specific adaptations to damage and environmental conditions.

## Main

Aging is a universal process in living organisms, and whether it is driven by a programmed genetic mechanism or by non-programmed, stochastic molecular damage remains a key area of debate in aging research^1, 2^. The programmed aging hypothesis proposes that aging is a genetically regulated process, similar to developmental stages such as growth and reproduction^3^. This theory suggests that evolutionary forces have selected mechanisms that actively promote aging to optimize population dynamics, facilitating generational turnover^4–6^. Supporting this view, genetic studies in model organisms such as zebrafish *Danio rerio* (*D. rerio*), fruit fly *Drosophila melanogaster* (*D. melanogaster*), and nematode *Caenorhabditis elegans* (*Nematode*) have demonstrated that modulating specific longevity pathways, including insulin/IGF-1 signaling (IIS), the target of rapamycin (TOR), and sirtuins, can significantly extend lifespan^7–9^. This evidence implies that aging may be controlled by evolutionarily conserved genetic programs, providing potential opportunities for manipulating aging through genetic or pharmacological interventions^10, 11^.

In contrast, the non-programmed aging hypothesis suggests that aging is not a predetermined biological program, but rather the result of cumulative molecular damage over time. Selective pressures favor traits that maximize reproductive success, placing less emphasis on somatic maintenance beyond reproductive age^12^. The free radical theory of aging, alongside other damage-based models, argues that aging is driven by oxidative stress, DNA damage, and protein misfolding, which gradually impair cellular function^13, 14^. From this perspective, although certain pathways may influence longevity, the aging process is primarily driven by damage accumulation. Interventions like caloric restriction and antioxidants are viewed as strategies to mitigate this damage rather than altering an intrinsic aging program^15, 16^.

Single-cell RNA sequencing (scRNA-seq) has rapidly advanced in recent years, allowing researchers to capture gene expression profiles of individual cells with unprecedented resolution^17, 18^. This technology enables the exploration of cellular heterogeneity and dynamic transcriptional changes across different tissues and developmental stages^19^. By analyzing scRNA-seq data, specific features can be identified to help distinguish between programmed and non-programmed processes. A programmed biological process should manifest in scRNA-seq as a stable co-expression module of related genes. If this process represents a fundamental, evolutionarily conserved biological function, the corresponding gene co-expression module should be consistently observed across different species and tissues^20^. Programmed processes would be expected to display co-expression modules that are conserved across organisms, particularly in genes related to essential functions such as stress response, metabolism, and protein homeostasis. Furthermore, such modules would be supported by conserved gene regulatory networks (GRNs) shared across species^21, 22^.

In contrast, a non-programmed biological process may not exhibit stable co-expression modules, even if orthologous genes are present across species. Instead, such processes may display species- or tissue-specific expression patterns, reflecting more stochastic or context-dependent regulation, rather than a conserved, genetically driven program. Processes driven by damage accumulation or environmental stress may lead to heterogeneous and asynchronous transcriptional activity, indicating the absence of tightly regulated GRNs.

In this study, we applied single-cell Weighted Gene Co-expression Network Analysis (WGCNA) to single-cell RNA sequencing data from zebrafish, fruit fly, and nematode across various life stages to identify age-related gene co-expression modules^23^. Our analysis identified a conserved gene module co-expressed during early life stages across all three species, specifically involved in the synthesis of various ribosomal proteins. In contrast, no shared co-expression module related to aging was detected across the organisms. Additionally, we constructed GRNs for each species^24^. By analyzing the GRNs underlying the co-expressed gene modules, we further demonstrated that there is no conserved, shared transcriptional regulatory network driving the aging process in these species. Based on these results, we suggest that there is no fundamental, ancient, cross-species programmed aging mechanism playing a dominant role in the aging process of different organisms.

## Results

### Tissue-Specific Age-Related Gene Modules in Zebrafish

A zebrafish single-cell sequencing dataset was downloaded from the Cross-Species Cell Landscape project^25^, which contains 1,082,680 cell samples spanning five distinct life stages of zebrafish, including 24 hours post-fertilization, the 72-hour larval stage, 21 days, 3 months, and 22 months, representing the aging stage (Fig. 1a). These cells were classified into 41 different cell types (Fig. 1b). For tissue- and age-specific analysis, we selected 11 cell types with higher cell counts to ensure robust data. These included endothelial cells (18,051), epithelial cells (104,543), intestinal bulb cells (57,032), macrophages (38,877), mesenchymal cells (89,466), muscle cells (13,432), neural cells (81,852), neural progenitor cells (72,417), osteoblasts (21,995), pancreatic cells (13,002), and primordial germ cells (19,443).

**Figure 1:**
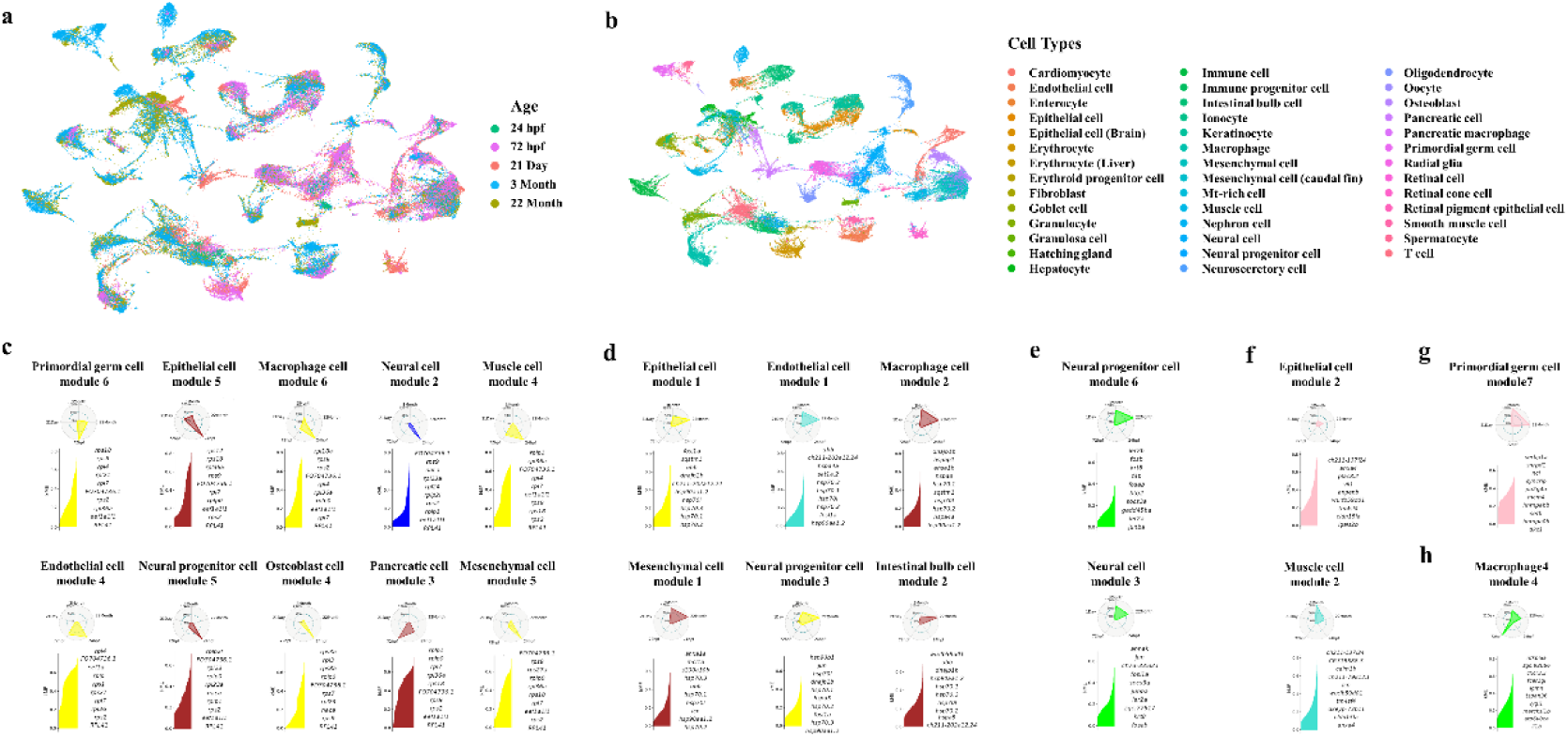
Identification of age-related gene modules across zebrafish life stages. a, UMAP plot of zebrafish cells across five life stages: This UMAP visualization displays the distribution of single cells from zebrafish sampled at five distinct life stages, including 24 hours post-fertilization (hpf), 72 hpf, 21 days, 3 months, and 22 months. Each point represents a single cell, colored by its respective time point, showing how cells cluster as zebrafish progress through developmental and aging stages. b, Cell type classification: This UMAP plot categorizes the cells into 41 unique zebrafish cell types. Each color corresponds to a different cell type, ranging from cardiomyocytes and immune cells to various progenitor and neuronal cells, providing a comprehensive view of the cellular landscape across zebrafish aging. c, Early-life expression of RPS/RPL modules in ten zebrafish tissues: This subfigure shows the expression of RPS/RPL modules (ribosomal protein genes) during early life stages in ten different zebrafish tissues. The top 10 genes with the highest kME (module eigengene connectivity) values are annotated for each tissue, highlighting key genes involved in ribosomal function during early development. d, Late-life expression of HSP-UPS modules in six zebrafish tissues: This subfigure highlights the HSP-UPS modules (genes involved in heat shock proteins and the ubiquitin-proteasome system) that are highly expressed during late life stages in six different zebrafish tissues. e, HSP-UPS modules in neural progenitor and neural cells: Cytoskeletal and immune stress-related genes are grouped into separate modules in neural progenitor cells (module 6) and neural cells (module 3) due to lower kME values, suggesting they are less central to these networks. f, Module 2 overlap in epithelial and muscle cells: Both epithelial cells (module 2) and muscle cells (module 2) share key hub genes involved in membrane stability and lipid metabolism, indicating independent regulation with minimal external influence. g, Primordial germ cell module 7: Primordial germ cell module 7 is an independent module, containing genes related to chromatin remodeling and ncRNA synthesis, aligning with aging hallmarks like genomic instability. h, Macrophage module 4: In macrophages (module 4), nfkbiaa, a regulator of the NF -κB pathway, is highly expressed, playing a key role in immune response and inflammation suppression during aging.

Then, we performed single cell WGCNA on the 11 selected cell types and identified a total of 46 age-related co-expression gene modules (Fig. S1a,b). These modules can be broadly divided into two categories based on their expression patterns. The first category is highly expressed during the juvenile stages, including module 4 of endothelial cells, module 5 of epithelial cells, module 6 of macrophages, module 5 of mesenchymal cells, module 4 of muscle cells, module 5 of neural progenitor cells, module 2 of neural cells, module 4 of osteoblasts, module 3 of pancreatic cells and module 6 of primordial germ cells (Fig. 1c). The genes within these modules are mainly related to ribosomal proteins and play key roles in protein synthesis. For instance, Ribosomal Protein S2 (RPS2), Ribosomal Protein S9 (RPS9), and Ribosomal Protein S18 (RPS18), components of the small ribosomal subunit, are involved in the initiation, elongation, and termination of mRNA translation. Ribosomal Protein L7 (RPL7) and Ribosomal Protein L23 (RPL23), part of the large ribosomal subunit, assist in peptide chain elongation and rRNA binding, contributing to ribosomal structure stability. Moreover, the *naca* gene encodes the nascent polypeptide-associated complex alpha subunit. Additionally, these genes are hub genes with high module membership (kME > 0.8) (Fig. 1c), suggesting they play a central role in the function of the first category of modules^26^. For convenience, we will refer to this module as the RPS/RPL module in the following text.

The second category of modules shows increased expression during adulthood and aging. In these modules, a variety of genes are shared across tissues, including those encoding proteins from the Heat Shock Protein (HSP) family, such as *hsp70* and *hsp90*, the ubiquitin B-encoding gene *ubb*, and genes encoding members of the Activator Protein-1 (AP-1) family, like *fosl1a* and *jun*, which encodes c-Jun, a component of the AP-1 complex^27^. These genes serve as hub genes (kME > 0.6) and present in several modules, including module 1 of epithelial cells, module 3 of neural progenitor cells, module 1 of endothelial cells, module 2 of macrophages, module 1 of mesenchymal cells, and module 2 of intestinal bulb cells (Fig. 1d). Proteins encoded by these genes are key components of pathways involved in protein misfolding, a process associated with aging and cellular stress^28^. HSP are induced in response to a variety of stressors related to aging and life span. It counteracts the cellular aging process by maintaining the structural and functional stability of paired proteins^29^. The protein encoded by the *ubb* gene is a crucial component of the ubiquitin-proteasome system (UPS), involved in key biological processes including cell signalling, cell cycle regulation, and apoptosis^30, 31^. It is also closely associated with the aging process across multiple species, from yeast to humans ^32^. Another key gene in this type of modules is *sqstm1*, which encodes Sequestosome 1 (SQSTM1), also known as p62. SQSTM1/p62 is a multifunctional protein that serves as a selective autophagy receptor, playing a key role in identifying and degrading damaged proteins or organelles. It binds to ubiquitinated proteins and directs them to autophagosomes for degradation^33^. In addition, SQSTM1/p62 contributes to the ubiquitin-proteasome pathway by binding to ubiquitin chains, facilitating the removal of misfolded or damaged proteins^34^.

Across the 46 modules analysed, HSP and UPS-related genes are present in 6 modules, distributed across 6 of the 11 zebrafish tissues, and consistently co-occur with genes encoding HSP family proteins, exhibiting high kME values. We refer to these as HSP-UPS modules, which display aging-specific expression. A distinct group of shared genes is found within the HSP-UPS modules across these 6 cell types (Fig. 1d). In addition to the HSP/UPS-related genes, these modules also comprise genes encoding cytoskeletal proteins, such as *ahnak* and *krt8*, as well as genes associated with immune stress responses, including *jun*, *fosl1a*, *socs3a*, *junba*, *ier2a*, and *fosab*, which are closely linked to aging. Cytoskeletal proteins are crucial for maintaining cellular structure and integrity, which can deteriorate with age, leading to cell dysfunction and tissue degradation^35, 36^. Meanwhile, genes associated with immune stress responses are involved in managing cellular damage and inflammation^37^. During the aging process, chronic low-level inflammation (inflammaging) and increased immune stress become more prevalent, contributing to the decline in tissue function and the onset of age-related diseases. In 4 of the 6 HSP-UPS modules from different cell types, nearly all genes encoding cytoskeletal proteins and immune stress-related genes appear simultaneously. However, in neural progenitor cells and neural cells, genes encoding cytoskeletal proteins are grouped into separate modules (module 6 of neural progenitor cells and module 3 of neural cells) due to lower kME values, which also suggest that they are subsidiary to other modules with higher kME values (Fig. 1e).

Apart from the HSP-UPS module, a distinct pattern is observed in module 2 of epithelial and module 2 of muscle cells. Although the epithelial cell module contains fewer genes, it completely overlaps with module 2 of muscle cells (Fig. 1f). Both modules contain high-kME hub genes (kME > 0.8), indicating that they are independently regulated with minimal influence from other biological processes. The shared hub genes include *anxa4*, *tm4sf4*, and *cldn15la*, all encoding membrane proteins. The gene *anxa4* encodes Annexin A4, a calcium-dependent phospholipid-binding protein involved in processes such as membrane trafficking, inflammation, and apoptosis^38, 39^. TM4SF4, a member of the tetraspanin family, plays a role in cell signalling, particularly in the regulation of cell migration and proliferation^40, 41^.

Additionally, module 2 of epithelial and module 2 of muscle cells include genes involved in lipid metabolism, such as *acbd7*, *acsl4a*, *cpt1ab*, *elovl1b*, and *pisd* (Table S1). They also find genes related to cell membrane formation and stability, such as *krt18*, *insig1*, and *pisd*, as well as stress-related genes like *ndrg1a*, *serpinb1*, and *sh3bgrl*. The *acbd7* gene encodes Acyl-CoA Binding Domain-Containing 7 (ACBD7), which is predominantly expressed in mammalian organs, including the brain, testis, and ovaries. Its homologs function as endogenous inhibitors of autophagy, regulating this process across fungi, plants, and animals^42^. Acyl-CoA Synthetase Long Chain Family Member 4a (ACSL4a), encoded by *acsl4a*, plays a key role in fatty acid metabolism. Recent studies indicate that the overexpression of *acsl4a* in zebrafish is triggered by Vitamin D receptor impairment, leading to hepatic steatosis ^43^. Meanwhile, *cpt1ab* plays a critical role in fatty acid oxidation (FAO)^44^, influencing aging through three mechanisms. First, FAO is vital for maintaining energy balance, particularly during fasting or caloric restriction, both of which has been linked to increased lifespan in various organisms. Second, as mitochondria age, impaired FAO leads to fatty acid accumulation and lipotoxicity, driving inflammation, tissue damage, and accelerating the aging process. Finally, FAO generates reactive oxygen species (ROS), which are closely associated with ROS-mediated aging processes^45^. Given this, we can align this module with established aging hallmarks and designate it the Transmembrane Protein-Dependent Energy-ROS Module, highlighting its role in energy metabolism and oxidative stress regulation^46^.

The analysis of primordial germ cell modules reveals that the age-specific module 7 has limited genes overlap with other modules, establishing it as a fully independent module. This module primarily includes genes involved in chromatin remodeling and ncRNA synthesis and maturation, such as *snrpd1*, *ncl*, *syncrip*, *mcm4*, *setb*, *hnrnpa0b*, and *dkc* (Table 1). We designate this as the Epigenetic and Genomic Instability Module, as the functions of these genes align closely with aging hallmarks, particularly epigenetic alterations and genomic instability.

**Table 1:**
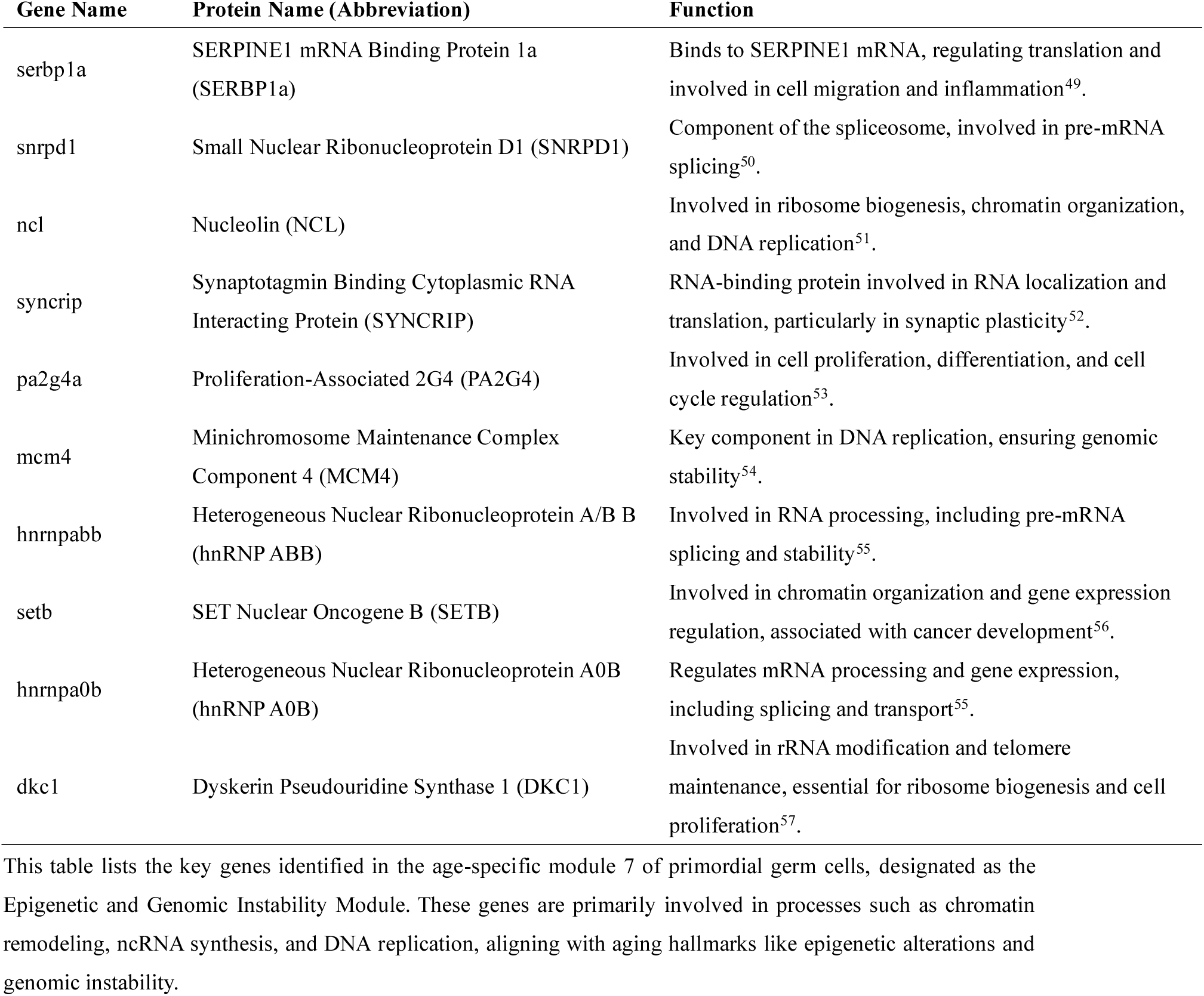
Key genes in the primordial germ cell epigenetic and genomic instability module.

In addition to the modules expressed only during early and late life stages, we observe module 4 in macrophages, with hub genes including nfkbiaa, which encodes NFKB Inhibitor Alpha A, an ortholog of the human version (Fig. 1h). This gene inhibits the NF-κB pathway, a key regulator of inflammation and immune responses^47^. Previous studies have demonstrated that suppressing the NF-κB pathway can delay the aging process in mouse models^48^. Unlike the previous two modules, the expression of this module in macrophages decreases between 3 and 22 months, consistent with its role in aging suppression. Notably, it is highly expressed at 72 hours post-fertilization, suggesting a potential involvement in cell differentiation and developmental processes. Based on the analysis, we designate module 4 of macrophages as the NFKB Inhibitory Module.

To confirm the conservation of these modules across different tissues, we created Venn diagrams using the top 30 genes with the highest kME values from the same type of modules in different tissues (Fig. 2a, b, c). We also performed KEGG and GO enrichment analyses on these modules and share gene between different module (Fig. 2d, e, f, g).

**Figure 2:**
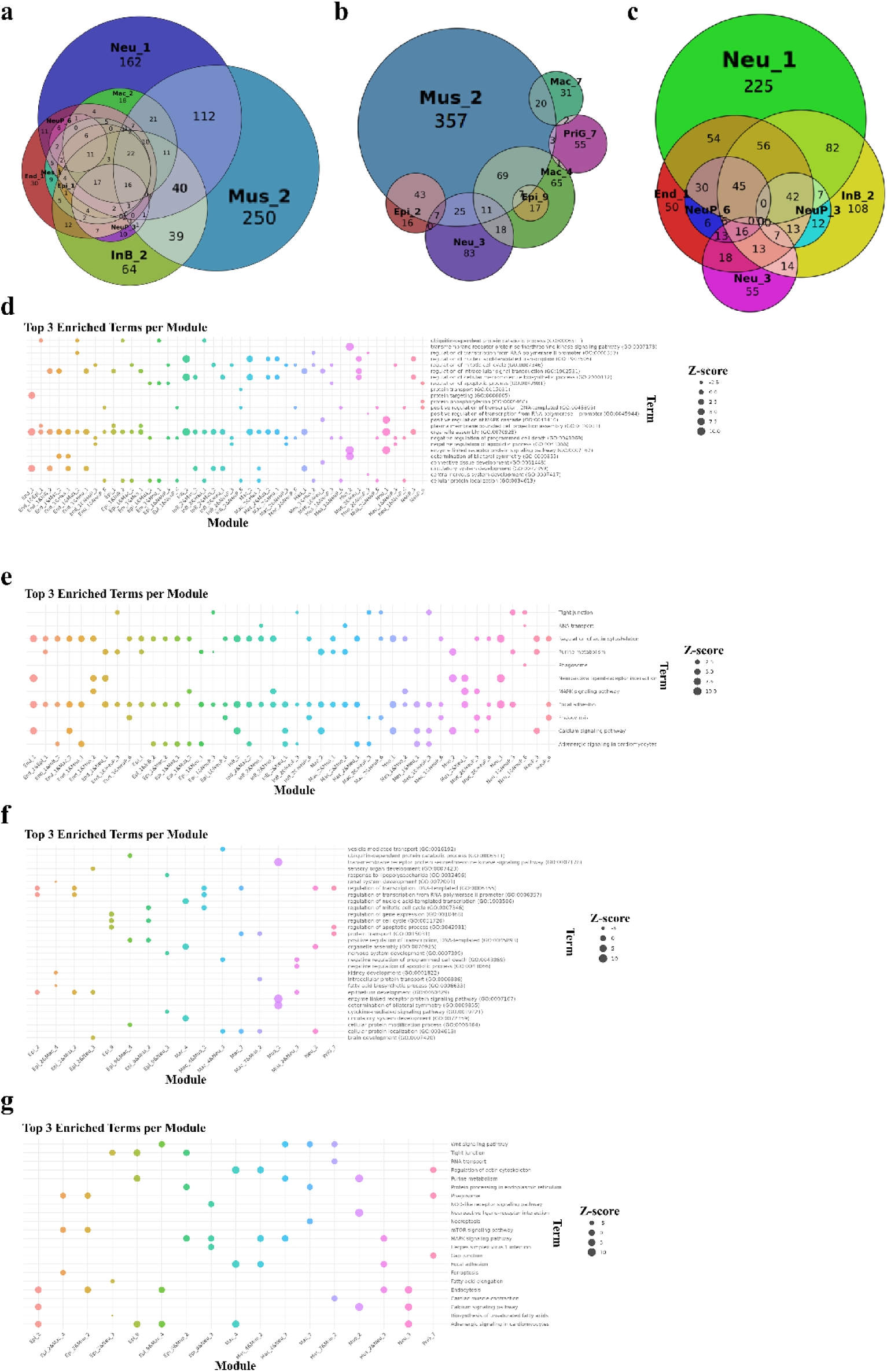
Conservation of gene co-expression modules across tissues and functional enrichment analysis. a-c, Venn diagrams depicting the overlap of the top 30 genes with the highest kME values across different tissues. These diagrams highlight shared and unique genes among modules of the same type across various tissues: a, shows modules from muscle cells, neurons, macrophages, and other tissues; b, focuses on muscle cell-specific modules; and c, presents overlapping genes in neural cell-specific modules. The numbers indicate the count of genes shared between or unique to specific modules. d-g, KEGG and GO enrichment analysis of the identified modules. Each dot represents the top 3 enriched biological processes or pathways per module. The Z-score (dot size) reflects the significance of the enrichment for each term. d, shows terms for modules related to neuronal development and immune function, e, focuses on cytoskeletal and signaling pathways, f, includes endocytosis and synaptic transmission pathways, and g, represents metabolic and cellular maintenance pathways.

In summary, through co-expression gene analysis of zebrafish at different ages, we identified four types of aging-related gene modules: the HSP-UPS Modules, the Transmembrane Protein-Dependent Energy-ROS Module, the NFKB Inhibitory Module, and the Epigenetic and Genomic Instability Module. These modules are linked to several key aging-related processes, including genomic instability, epigenetic alterations, loss of proteostasis, deregulated nutrient sensing, and mitochondrial dysfunction. Additionally, their selective expression across different cell types aligns with recent findings that aging processes are diverse and tissue specific.

### Tissue-Specific Aging-Related Gene Modules of Fruit fly

We utilized single-cell transcriptomic data of *D. melanogaster* from the Aging Fly Cell Atlas, published in 2023. This project sequenced both head and body tissues across four age stages (5, 30, 50, and 70 days), yielding over 868,000 nuclei, encompassing 17 cell-type categories and 163 subtypes (Fig. 3a, b) ^58^. From this vast dataset, we selected 10 cell types distributed across various tissues to identify and compare age-specific expression modules in fruit fly. These included olfactory receptor neurons (5,415 cells), outer photoreceptors (21,239 cells), R8 photoreceptors (3,108 cells), head skeletal muscle cells (7,765 cells), epithelial cell (44094), follicle cell (26610), crop (7047), germline cell (3824), indirect flight muscle (18057) and muscle cell (54592).

**Figure 3:**
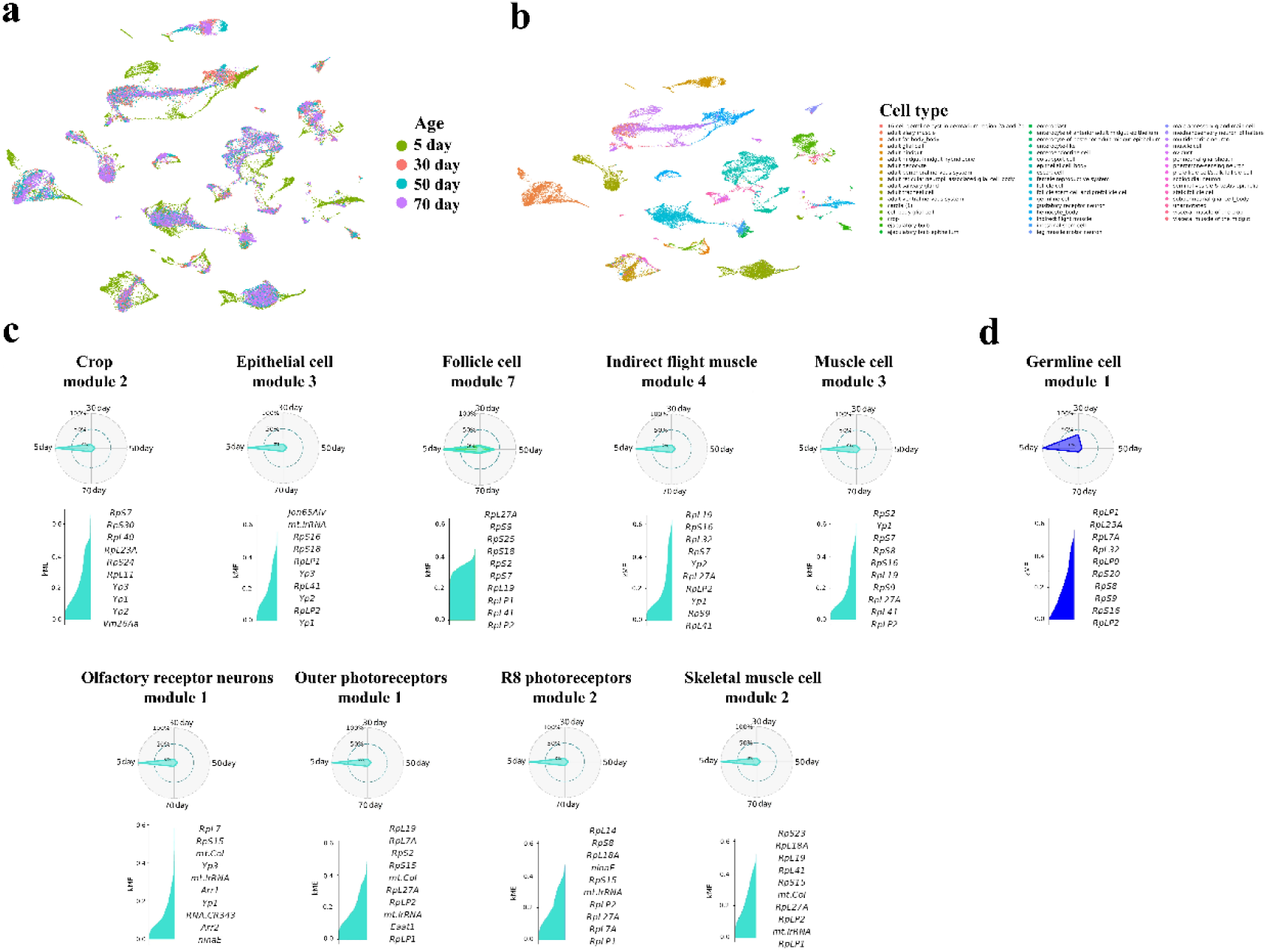
Analysis of fruit fly head and body tissues across four age stages. a, UMAP plot of nuclei from head and body tissues: The UMAP plot displays over 868,000 nuclei across four distinct age stages: 5, 30, 50, and 70 days. Each point represents a nucleus, and the colors indicate the different age groups, showing the distribution of cells during development and aging. b,Classification of 17 cell types and 163 subtypes: The UMAP plot categorizes nuclei into 17 major cell types and 163 subtypes. These include diverse cell categories such as muscle, nervous system, and germline cells, allowing for an in-depth analysis of tissue-specific expression changes throughout aging. c, Early expression RPS/RPL modules in various cell types: This panel highlights the conserved RPS/RPL gene co-expression modules found in 10 selected cell types, predominantly active during early developmental stages. The identified hub genes encode ribosomal proteins and other essential proteins like Jon65aiv in epithelial cells, involved in Johnston’s Organ Neuron function, and tropomyosin 2 (tm2) and neurexin IV (nrm), which are associated with muscle and nervous system development. d, Germline-specific early expression module: Unlike other cell types, germline cells maintain an active early transcription module encoding ribosomal proteins (RPS/RPL) even at 30 days of age, indicating prolonged expression compared to other tissues. This module contains key ribosomal protein genes, including mt.lrrna.

In our analysis, we discovered a conserved early expression RPS/RPL module present in 10 of the 10 selected fruit fly cell types, predominantly active during the early stages of cell development. Notably, germline cells also exhibited an early transcription module encoding ribosomal proteins (Fig. 3c). However, unlike other cell types, this module remained active in germline cells even at 30 days of age (Fig. 3d). The hub genes of this module included those encoding ribosomal proteins, such as mitochondrial ribosomal RNA (*mt.lrrna*), along with genes related to cell differentiation, like *jon65aiv* in epithelial cells, which encodes a protein associated with Johnston’s Organ Neuron^59^. Additionally, early expression modules associated with muscle and nervous system development were identified in indirect flight muscle cells and muscle cells. These modules involved genes such as tropomyosin 2 (*tm2*), neurexin IV (*nrm*), muscle LIM protein 60a (*mlp60a*), WD repeat domain 62 (*wdr62*), and neuroligin 1 (*nlg1*).

In contrast to the conserved early expression modules, late-life expression modules in fruit fly exhibited greater diversity in gene expression (Fig. 4a). Gene Ontology (GO) enrichment analysis of the top 30 genes (ranked by kME) from these modules revealed significant enrichment in transcriptional regulation, including terms such as DNA-templated transcription (GO:0006355) and regulation of transcription from RNA polymerase II promoter (GO:0006357) (Fig. 4b). These findings suggest that transcriptional regulation remains highly active in late life across fruit fly tissues, but these regulatory processes are not significantly enriched in typical stress and damage response pathways or related biological processes. This implies that the heightened RNA transcriptional regulation observed during fruit fly aging may represent an independent aging phenomenon, distinct from other common hallmarks and mechanisms of aging. Additionally, we observed widespread ectopic expression of certain genes across tissues, such as *norpA* (typically associated with photoreceptors) and *rut* (involved in learning and memory)^60, 61^, which were unexpectedly expressed in crop cells (digestive cells). Similarly, the gene *Sap47*, which encodes a protein involved in regulating synaptic function, and *tomosyn*, associated with synaptic vesicle release, are normally expressed in neurons of fruit fly. However, during the late stages of the fruit fly’s lifespan, these genes exhibited ectopic expression in epithelial cells ^62^. Furthermore, genes such as *fra* (involved in axon guidance) and *eya* (a transcription factor associated with eye development) were ectopically expressed in follicle cells of aging flies. Similar phenomena of ectopic gene expression during aging have also been documented in humans, where disrupted epigenetic controls lead to the activation of inappropriate lineage-specific genes^63^.

**Figure 4:**
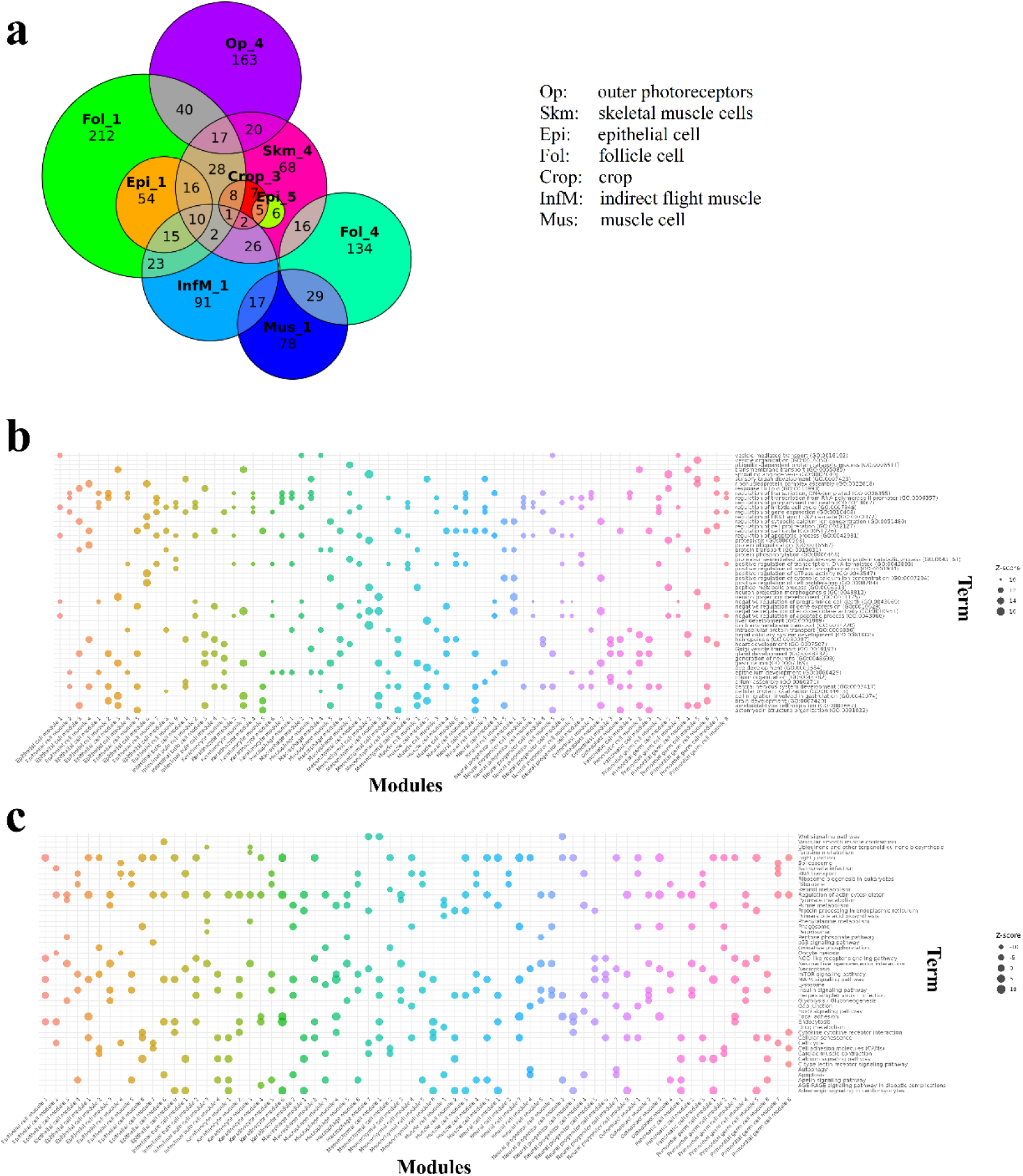
Late-life expression diversity and pathway enrichment in fruit fly tissues. a, Venn diagram of late-life gene expression modules across tissues: This diagram illustrates the diversity of gene expression modules in late-life stages of fruit fly across various tissues, including outer photoreceptors, skeletal muscle, epithelial cells, follicle cells, crop cells, and indirect flight muscle. The overlap between different tissue modules is shown, highlighting shared and unique genes in each tissue. b, GO enrichment analysis of transcriptional regulation in late-life modules: Gene Ontology (GO) analysis of the top 30 genes from each module reveals significant enrichment in terms related to transcriptional regulation, such as DNA-templated transcription and regulation of transcription from RNA polymerase II promoter, indicating that transcriptional regulation remains highly active across fruit fly tissues in late life. c, KEGG pathway enrichment analysis of aging-related pathways: Pathway enrichment analysis identifies key pathways involved in aging, particularly MAPK, mTOR, and Hippo signaling pathways, across different tissues. These pathways are significantly enriched in muscle and epithelial cells, reflecting their roles in regulating oxidative stress responses, nutrient sensing, and cellular repair mechanisms during aging.

A distinctive feature of late-life expression modules in fruit flies is the prevalence of non-coding RNAs (ncRNAs), particularly long non-coding RNAs (lncRNAs) and antisense RNAs (asRNAs). In germline cell module 3, ncRNAs comprised the majority of high-kME genes (7 out of 10), underscoring their increasing importance as epigenetic regulators that can potentially influence the aging process^64, 65^. In addition, members of the Argonaute protein family, *ago1* and *ago3*, were also observed in fruit fly aging modules across all nine cell types. Interestingly, *ago1* was exclusively expressed in epithelial cell (module 5), follicle cell (module 2), and indirect flight muscle cell (module 5), while *ago3* was confined to crop cell (module 3), germline cell (module 3), muscle cell (module 1), olfactory receptor neuron (module 5), outer photoreceptor cell (module 4), and skeletal muscle (module 3). This suggests a mutually exclusive expression pattern for *ago1* and *ago3*. The *ago1* is primarily involved in the miRNA pathway, where it binds to miRNAs to mediate the silencing of target mRNAs. miRNAs are small non-coding RNAs that guide Argonaute proteins to specific mRNA targets, resulting in either degradation or inhibition of translation. Argonaute proteins, including Ago1, are essential components of the RNA-induced silencing complex (RISC) and play a key role in RNA interference (RNAi) and post-transcriptional gene silencing mechanisms^66, 67^. Meanwhile, Ago3 is crucial in small RNA regulatory pathways, particularly in the PIWI-interacting RNA (piRNA) pathway, where it helps maintain genomic integrity in germ cells^68^. In *D. melanogaster*, Ago3 specifically targets transposable elements, preventing their integration into new DNA locations, which could otherwise cause genomic instability. This mechanism is essential for preserving germline stability and fertility. Ago3 works in conjunction with Aubergine (another Argonaute protein) to mediate this defence mechanism, ensuring proper germline function in fruit flies.

We conducted KEGG pathway enrichment analysis on the genes within these modules (Fig. 4c), identifying key aging-related pathways, particularly the MAPK, mTOR, and Hippo signalling pathways. The MAPK pathway was significantly enriched in muscle and epithelial cells, indicating increased oxidative stress responses and the need for enhanced cellular repair mechanisms as aging progresses^69^. This pathway’s dual roles in cell proliferation and apoptosis highlight its importance in maintaining tissue integrity, particularly in aging muscle and epithelial tissues, where it regulates both repair and survival processes. Additionally, the mTOR signalling pathway was enriched in crop, epithelial, and muscle cells, suggesting disruptions in nutrient sensing and metabolic regulation during aging^70^. The mTOR pathway regulates protein synthesis and autophagy, essential for maintaining muscle mass and metabolic balance, and its dysregulation is a hallmark of aging, contributing to muscle atrophy and metabolic inefficiency. Finally, the Hippo signalling pathway, highlighted in follicle and muscle cells, regulates cell growth, tissue homeostasis, and stem cell function^71^. Dysregulation of Hippo signalling is linked to age-related tissue degeneration and impaired tissue repair. In neural tissues, it likely contributes to neuroprotection and influences neurodegenerative processes during aging.

In conclusion, our comprehensive analysis of aging modules in fruit fly tissues reveals that transcriptional dysregulation is a hallmark of aging, characterized by the widespread activation of inappropriate transcriptional programs across various tissues. Furthermore, classical signalling pathways, including MAPK, mTOR, and Hippo, play significant roles in aging across different tissues, contributing to the regulation of cellular stress responses, metabolic shifts, and tissue homeostasis.

### Tissue-Specific Aging-Related Gene Modules in Nematodes

We utilized the single-cell transcriptomic data published by Roux, A.E. et al. in 2023, which covers the full life cycle of *Nematode*^72^. This dataset was collected at six distinct time points during *Nematode* adulthood (days 1, 3, 5, 8, 11, and 15). To enrich for somatic cells, temperature-sensitive gon-2 (q388ts) mutants were used, effectively reducing the number of germ cells^72^. The dataset consists of 211 distinct cell types, categorized into 14 major groups. From these, 8 cell types were selected for WGCNA analysis: epithelial cells (11,199), germ cells (983), glial cells (1,871), interneurons (6,342), motor neurons (8,966), muscle cells (3,938), general neurons (18,683), and seam cells (2,363). General neurons represent a mixture of neuron types, excluding motor, interneurons, and sensory neurons. A total of 74 co-expression modules were identified across these cell types.

In line with previous findings in zebrafish and fruit flies, we identified co-expression modules containing ribosomal protein-encoding genes (from the *rps* and *rpl* gene families) in 7 of the 8 selected cell types (Fig. 5a), with kME values greater than 0.6. These are referred to as RPS/RPL modules, primarily comprising genes involved in ribosomal functions, critical for protein synthesis and cellular maintenance. Additionally, in epithelial cells, interneurons, motor neurons, muscle cells, and seam cells, two genes with currently unclear functions, *W01D2.1* and *Y37E3.8*, were co-expressed within the RPS/RPL module, indicating potential roles in early cell development. Furthermore, unlike in zebrafish and fruit flies, the expression of the RPS/RPL module in nematode is not limited to early life stages, it remains expressed during the late stages (day 15) of life in epithelial cells, germ cells, and neurons.

**Figure 5:**
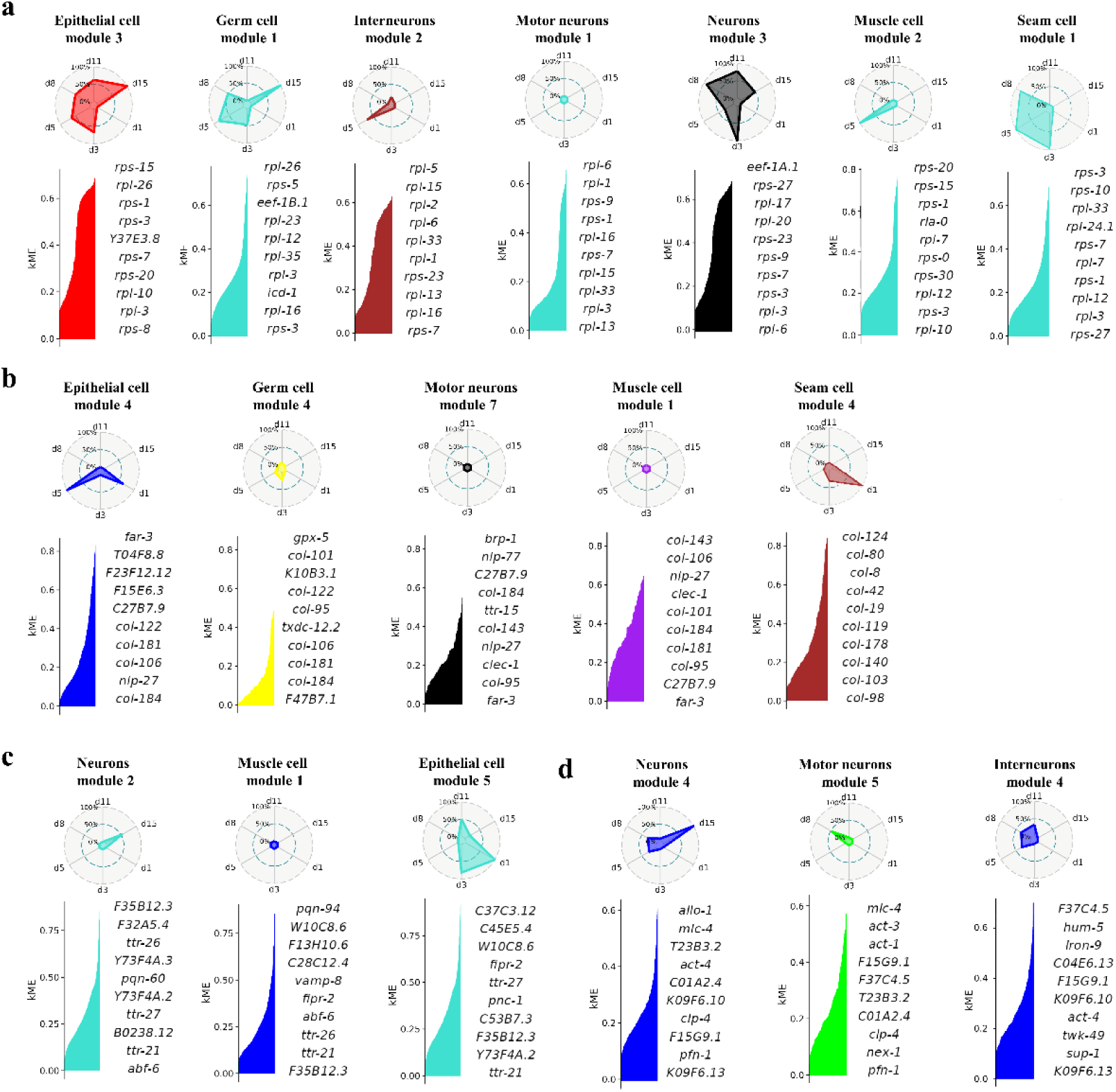
Co-expression modules in nematode across cell types and developmental stages. a, RPS/RPL modules in various nematode cell types: Co-expression modules composed of ribosomal protein-encoding genes (*rps* and *rpl*) were identified in 7 out of 8 selected cell types, including epithelial cells, interneurons, motor neurons, muscle cells, and seam cells. b, Col-nlp modules in epithelial, germ, and seam cells: This panel shows the early-life co-expression module named the col-nlp module, primarily composed of collagen (*col*) and neuropeptide-like protein (*nlp*) family genes. c, Co-expression modules in neurons, muscle cells, and epithelial cells in nematode. The genes are *ttr-26*, *ttr-27*, and *ttr-21*. d, Profilin and actin-related modules in neuronal cell types: In general neurons, motor neurons, and interneurons, a co-expression module consisting of *pfn-1, act-1, act-4, did-2*, *nex-1*, and *clp-4* was identified.

In *Nematode* epithelial cells, germ cells, and seam cells, we identified an early-life co-expression gene module, which we have named as col-nlp module for the purposes of this study (Fig. 5b). This module is primarily composed of genes encoding the collagen family (*col*) and the neuropeptide-like protein family (*nlp*). The *col* genes are predicted to encode collagen proteins involved in cuticle formation, a major structural component of nematode^73^. Collagen is crucial for maintaining cuticle integrity, helping the organism resist environmental stress and maintain mobility. The neuropeptide-like proteins encoded by *nlp-29* and *nlp-33* play an important role in immune responses and stress signaling^74–76^. Additionally, this module includes the *far-3* gene, which encodes a Fatty Acid and Retinol-binding Protein (FAR family). This protein binds to fatty acids and retinoids (derivatives of Vitamin A), participating in lipid metabolism and fatty acid transport. In nematode, *far-3* is likely involved in lipid storage, energy metabolism, and cell signalling. These proteins help regulate energy metabolism and maintain cell membrane integrity, particularly under environmental stress^77^. The genes in this module are often upregulated in response to infection or stress, helping nematode to defend against pathogens and manage stress. Notably, this module is identified as silent across the lifecycle in motor neurons and muscle cells.

The neurons (module 2), muscle cells (module 1), and epithelial cells (module 5) in Nematode highlight several key genes involved in critical cellular functions (Fig. 5c). Among these genes, *ttr-26, ttr-27*, and *ttr-21* encode transthyretin-like proteins, which are implicated in the binding and transport of small molecules, potentially playing a significant role in hormone regulation and protein stability. These proteins are essential for maintaining cellular homeostasis, particularly under stress conditions. The expression profiles of these genes—both shared across tissues and specific to certain cell types—underscore the critical roles of TTR-like proteins, vesicle transport mechanisms, and immune-related responses in Nematode development and stress management. The identified genes suggest a coordinated inter-tissue effort to ensure protein stability, energy metabolism, and cellular resilience under various stressors, reflecting the organism’s ability to manage environmental and physiological challenges.

In the three selected neuronal cell types, including general neurons, motor neurons, and interneurons, we identified a co-expression module consisting of *pfn-1*, *act-1*, *act-4*, *did-2*, *nex-1*, and *clp-4*. The *pfn-1* gene encodes Profilin, a crucial regulator of actin polymerization involved in key processes such as cell movement, division, and the maintenance of cellular structure^78^. Genes act-1 and act-4 encode Actin, which participates in vital processes such as maintaining cell structure, movement, and muscle contraction^79^. Meanwhile, genes did-2 and nex-1 are involved in vesicle-mediated transport and degradation of cellular components^80, 81^. The co-expression of these genes suggests that the cells may be undergoing processes such as mitosis and vesicle transport. It is important to note that the timing of this module’s expression differs among the three neuron types (Fig. 5d). In general neurons, the module is expressed on days 5, 8, and 11, with high expression on day 15. In motor neurons, the module is only expressed on day 8, while in interneurons, it is expressed on days 5, 8, and 11. This result demonstrates that our method can identify the asynchronous development characteristics of similar cell types in *Nematode*, which is a defining feature of nematode development ^82^.

In addition to the aforementioned co-expression modules, genes encoding the HSP superfamily exhibit variations in expression levels at different time points across the eight cell types studied. However, unlike in zebrafish, these genes do not form a unified co-expression module. The genes include *hsp-1*, *hsp-3*, *hsp-4*, *hsp-6*, *hsp-16*, *hsp-25*, *hsp-43*, *hsp-70*, *hsp-90*, and *hsp-110*. Unlike zebrafish, where *hsp* gene expression tends to increase with age, the expression patterns of *hsp* genes in nematode show no consistent age-related trend and vary significantly across different cell types (Fig. 6). For instance, in epithelial cells, *hsp-70* consistently shows low expression, while *hsp-1* and *hsp-90* peak at day 15. Other *hsp* genes are not expressed at all on day 15. In glial cells, interneurons, motor neurons, and seam cells, *hsp* gene expression is lowest on day 15. In muscle cells, all *hsp* genes reach their lowest expression on day 11, and this level persists through day 15. By contrast, in germ cells, *hsp-70* and *hsp-90* reach their highest expression levels on day 15, similar to *hsp-90* in general neurons. When compared to the zebrafish HSP-UPS module, two features of *hsp* gene expression in nematode become apparent: (1) the various *hsp* genes in nematode do not display co-expression, indicating the absence of a unified transcriptional regulatory mechanism governing *hsp* expression; (2) the timing of *hsp* expression increases varies across cell types, with a substantial number of cells shutting down *hsp* gene expression in late life stages.

**Figure 6:**
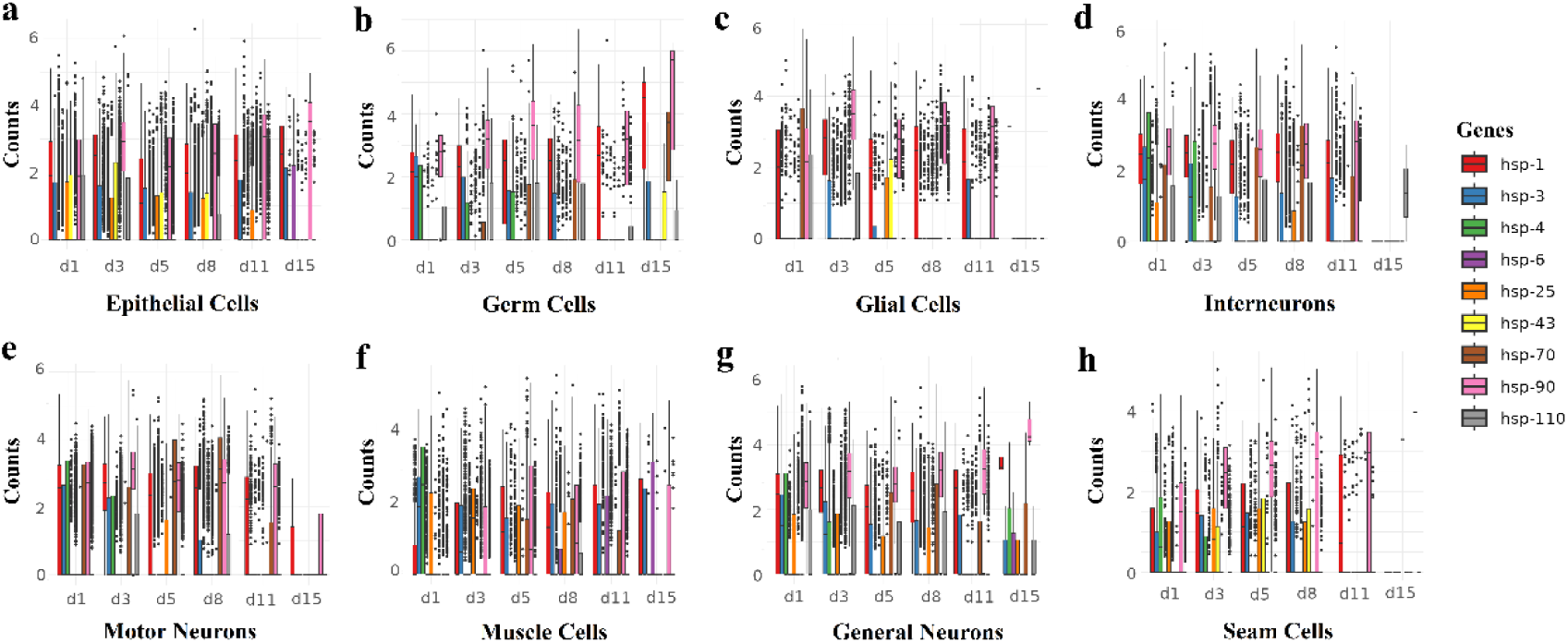
Temporal expression patterns of HSP genes across eight *C. elegans* cell types. a, In epithelial cells, the expression of *hsp-70* remains consistently low, while *hsp-1* and *hsp-90* peak at day 15, suggesting a late-life increase in stress response. Other *hsp* genes show variable expression, with some not expressed at all on day 15. b, Germ cells: Both *hsp-70* and *hsp-90* exhibit the highest expression at day 15. The remaining *hsp* genes are less consistently expressed over time. c, Glial cells: *hsp* gene expression is generally lower on day 15 compared to earlier time points. d, Interneurons: The *hsp* gene expression declines by day 1. e, Motor neurons:*hsp-1* and *hsp-90*, show reduced expression by day 15. f, Muscle cells: All *hsp* genes exhibit their lowest expression levels by day 11, with this reduction persisting through day 15. g, General neurons: *hsp-90* expression increases by day 15. h, Seam cells: *hsp* genes expression is lowest on day 15 in seam cells.

In summary, co-expression gene analysis of *Nematode* reveals that different cell types follow distinct life cycles and enter the aging process at different times. This finding aligns with previous studies that have shown variability in the aging onset across different tissues in nematode^83^. A significant portion of cells during the aging phase downregulate the expression of genes associated with repair and maintenance of cellular homeostasis, effectively ceasing their protective responses against the aging process.

### Age-Related Changes in Gene Expression Regulation

To investigate whether the transcriptional regulatory mechanisms underlying co-expressed gene modules differ between species, we constructed gene regulatory network (GRN) models for zebrafish, nematode and fruit fly across all tissues. We extracted the subnetwork containing the nine genes with the highest average kME values (kME > 0.7) from the late-stage co-expression module of heat shock proteins (HSPs) and ubiquitin-proteasome system (UPS) in zebrafish, forming the zebrafish HSP-UPS local GRN (Fig. 7a). This subnetwork introduced a total of 14 transcription factors, 13 of which were additional and one of which, *fosl1a*, was already present among the 9 high-kME genes. We then identified the homologous genes of these 22 genes in fruit fly and nematode (Table S2) and built corresponding HSP-UPS GRNs for each species (Fig. 7b,c).

**Figure 7:**
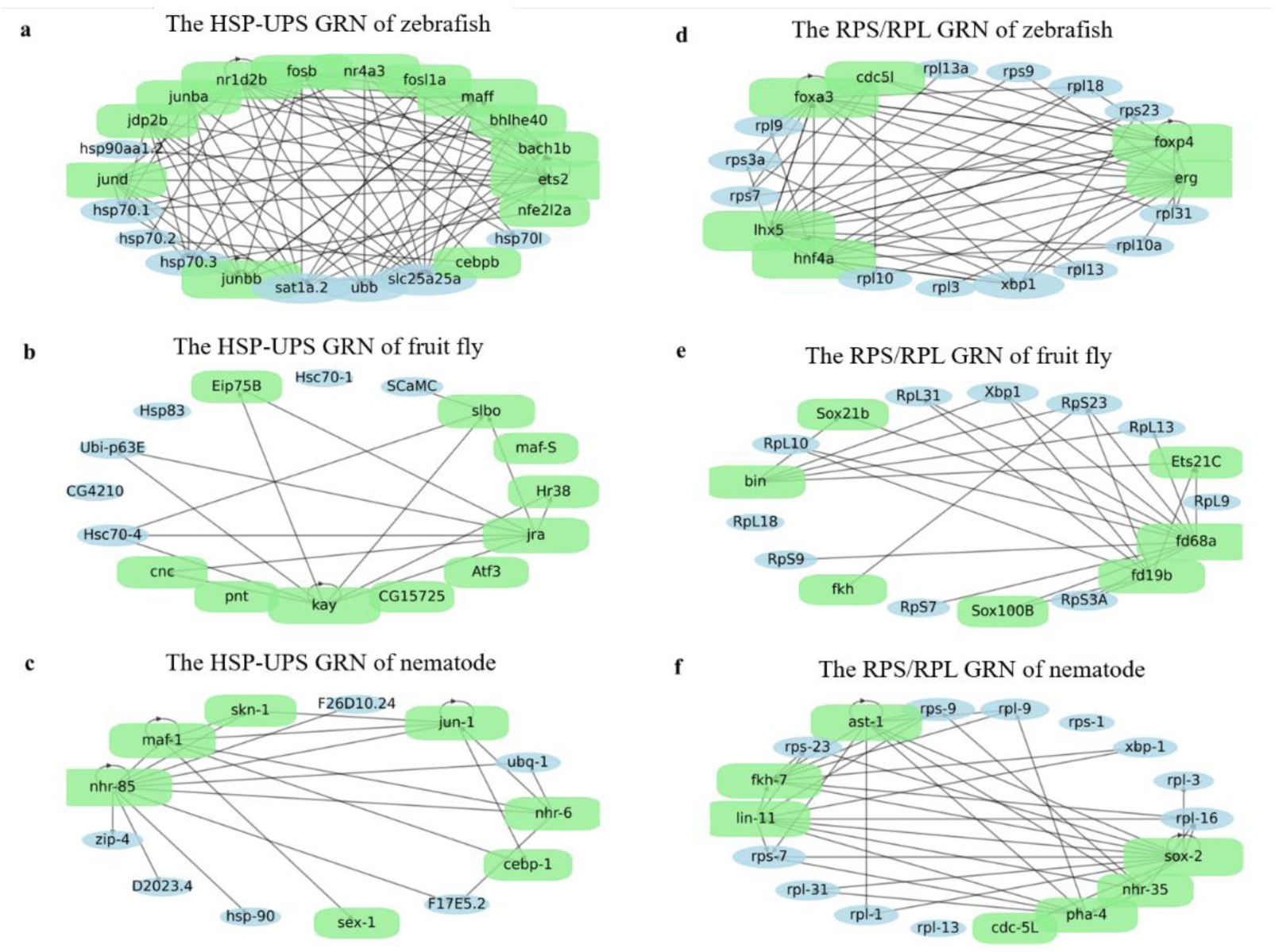
Gene Regulatory Networks (GRNs) of HSP-UPS and RPS/RPL modules in zebrafish, fruit fly, and nematode. a, The HSP-UPS GRN in zebrafish displays a complex regulatory network with transcription factors such as *ets2*, *nr1d2b*, and *fosl1a* orchestrating the expression of key HSP genes like *hsp90a1.2*, *hsp70.1*, and *hsp70.2*. b, In the fruit fly, the HSP-UPS network is less complex, with fewer transcription factors, including *Jra* and *pnt*, regulating *Hsc70-4* and *Ubi-p63E*. c, The nematode HSP-UPS GRN also shows a distinct regulatory architecture. Transcription factors *nhr-85* and *nhr-6* regulate HSP genes like *hsp-90* and *ubq-1*. d, The RPS/RPL GRN in zebrafish reveals transcription factors such as *foxa3*, *foxp*, and *hnf4a* tightly regulating ribosomal protein genes like *rpl13a* and *rps9*. e, In the fruit fly, FOX family genes like *fkh* and *fd68A* play major roles in regulating ribosomal proteins such as *Rpl31* and *Rps23*. f, The nematode RPS/RPL GRN includes transcription factors like *sox-2*, *nhr-35*, and *fkh-7*, which regulate the expression of ribosomal genes such as *rps-9* and *rpl-13*.

Through analysis of the HSP gene subnetworks, we observed tight transcriptional regulation between HSP genes, such as *hsp70* and *hsp90*, and transcription factors in zebrafish. In zebrafish, the highest degrees of connectivity within the HSP-UPS network came from *ets2*, *nr1d2b*, and members of the *jun* family, which mediated the expression of most *hsp* family genes. Genes *ets2*, *nr1d2b* (encoding the nuclear receptor subfamily 1 group D member 2), and the *Jun* family (orthologous to the human *Jun proto-oncogene*, a subunit of the AP-1 transcription factor) are predicted to act as transcription factors with DNA-binding activity and RNA polymerase II specificity. In the fruit fly HSP-UPS GRN (Fig. 7b), the complexity of this network was reduced, with an average degree of 0.5, significantly lower than zebrafish (average degree of 6) and nematode (average degree of 3.5) HSP-UPS GRNs. In the fruit fly GRN, *Jun* family transcription factors like *Jra* regulated *Ubi-p63E* and *Hsc70-4*, similar to their roles in zebrafish and nematodes. However, the expression of *Jra* was very low in the single-cell transcriptomics data of nematodes, and it was absent in all identified co-expression modules. In nematodes (Fig. 7c), the transcription factors *nhr-85* and *nhr-6*, orthologs of zebrafish *nr1d2b* and *nr4a3*, exhibited similar regulatory functions. These transcription factors regulated the expression of *hsp* family genes (*hsp90* and *F26D10.24* in nematodes) and the ubiquitin ligase-encoding genes *ubb/ubq-1*, suggesting conservation of this regulatory pathway across species. However, the nematode *jun-1* gene, a direct ortholog of the *jun* family, did not exhibit similar regulatory functions in the nematode GRN. Comparing the HSP-UPS GRNs of these three model organisms reveals that HSP-UPS regulatory patterns are not conserved across species. Although HSP-UPS genes are widely present across species, their transcriptional regulatory mechanisms differ significantly.

As a comparison, we also analysed the local GRNs of the RPS early-expression modules with more similar co-expression characteristics in the three species to assess the transcriptional regulatory relationships involved in a well-defined, conserved biological process. In zebrafish, six transcription factors co-regulated the expression of 13 ribosomal protein-coding genes in the GRN (Fig. 7d). These transcription factors included *foxa3* and *foxp*, which encode members of the forkhead box (FOX) family, *cdc5l* (encoding cell division cycle 5-like protein), *lhx5* (orthologous to human *LHX5*, a LIM homeobox protein), *hnf4a* (orthologous to human *HNF4A*, hepatocyte nuclear factor 4 alpha), and *erg* (orthologous to human *ERG*, ETS transcription factor ERG). In the fruit fly RPS early-expression module GRN (Fig. 7e), FOX family genes such as *fkh* (FoxA), *fd68A* (FoxK), *fd19B* (FoxG), and *bin* (FoxF) directly regulated all *rpl* and *rps* genes. The widespread expression and multifunctionality of FOX family genes, which span functions from cell fate determination to organ-specific development, underline their crucial role in embryonic development and the maintenance of tissue homeostasis in adult organisms. In nematodes, five out of the six transcription factors from the zebrafish RPS/RPL GRN (except *cdc5l*) had nematode orthologs (Fig. 7f), including *sox-2*, *nhr-35*, *lin-11*, *fkh-7*, *ast-1*, and *pha-4*, regulating the expression of *rps/rpl* genes. Among them, *sox-2*, the *Nematode* ortholog of Sox2, is a key transcription factor involved in cell fate determination, embryonic development, and stem cell maintenance, and its conservation across the three species underscores the evolutionary importance of its regulatory function^84^. Another important transcription factor, *ast-1*, is the direct ortholog of zebrafish *erg* and plays a critical role in angiogenesis and cell proliferation, influencing processes such as cell division and blood cell formation^85^.

In terms of network connectivity, zebrafish showed the highest average node degree (15), suggesting a robust regulatory relationship between transcription factors and target genes, with high network complexity. The fruit fly GRN had an average node degree of 10, while the nematode GRN fell in between at approximately 12. Comparing the RPS/RPL GRNs of the three species, it is evident that the nematode GRN more closely resembles that of zebrafish, consistent with the observations from the HSP-UPS GRN. The smaller difference in network complexity among the RPS/RPL GRNs across the three species compared to their HSP-UPS GRNs further indicates that the RPS/RPL network is more evolutionarily conserved across species. This greater conservation in the RPS/RPL GRN suggests that transcriptional regulation of ribosomal protein genes plays a more fundamental and consistent role in cellular processes across diverse organisms than the regulation of heat shock proteins.

Thus, the comparison of RPS/RPL gene regulatory networks among zebrafish, fruit fly and nematode reveals that transcriptional regulation related to ribosomal proteins is highly conserved. In contrast, the HSP-UPS GRN shows greater species-specific variations. This supports the hypothesis that while genes involved in stress responses like *hsps* are shared across species, their regulation may be more adaptable to specific environmental or physiological contexts, whereas the transcriptional regulation of fundamental cellular processes such as protein synthesis tends to be highly conserved.

## Discussion

The debate over whether aging is a programmed or non-programmed process has long been rooted in evolutionary theory and hypotheses derived from ecological models. Programmed aging hypotheses propose that aging might serve as an evolutionarily advantageous mechanism for population turnover, while non-programmed theories suggest that aging results from the accumulation of damage and errors throughout an organism’s life. However, both hypotheses lack direct empirical evidence from experimental biology and have yet to be substantiated through detailed molecular and genetic studies.

In this study, we addressed this question by analysing single-cell RNA sequencing data from zebrafish, fruit fly, and nematode across different life stages. Our findings demonstrate the absence of a shared, evolutionarily conserved programmed mechanism underlying aging across these species. After comparison, we observed that a truly programmed process, the transcription of ribosomal protein genes RPS/RPL, reflected by gene co-expression in single-cell transcriptomic data, is conserved across species during early developmental stages. In all three species, the RPS/RPL module displays conservation not only in gene composition but also in its expression pattern, reinforcing the notion that early-life developmental programs are tightly regulated and evolutionarily conserved, even among highly divergent organisms^86^. The identification of the early-expressed RPS/RPL co-expression module confirms that our approach can effectively identify programmed biological processes.

By contrast, the gene expression modules involved in aging are not only highly diverse across different species but also exhibit significant variability across different tissues within the same organism. The widespread co-expression of HSP-related genes in somatic cells of aging zebrafish was not detected in the germ cells of aging zebrafish, nor was it found in aging cells of fruit flies and nematodes. The aging cells of fruit flies exhibited active transcription, with diverse ectopic gene expression reflecting a loss of transcriptional control during aging. In nematodes, the diversity in the aging process also includes differences in the onset timing of aging in different cells; for example, the co-expression module related to act genes in different neuron types of nematodes shows varied transcriptional timing. Our findings align with previous studies of tissue-specific aging across species, particularly those emphasizing the heterogeneity of aging. For example, single-cell RNA sequencing of aging in mouse kidneys, lungs, and spleens revealed that distinct cell types express unique aging-related genes, each following different enrichment trajectories^87^. In humans, single-cell transcriptomic analyses of oocytes have highlighted the differentiation of gene expression related to DNA damage and apoptosis^88^, while in the aging human cortex, immune-related HLA-DPA1 and protein folding-related genes in the HSP40 family show the greatest expression changes^89^. The latest single cell sequencing results of human brain aging indicate that the vast majority of differentially expressed (DE) genes were unique to a single cell type^90, 91^. This tissue-specific diversity in aging mechanisms further supports the idea that aging is not governed by a conserved, universal program, and reinforces the notion that there is no single, universally conserved programmed aging process.

The recently proposed Cell Aging Regulation System (CARS) theory describes aging as a system that integrates various signals and relies on the nuclear genetic Aging Program to determine how cells age^92^. However, our results raise questions about the universality of such a system. If CARS is entirely spontaneous and forms independently of cellular damage pressure, why doesn’t it utilize the same gene products as internal aging signals across tissues and species? Specifically, while we identified high expression of HSP-related genes in aging zebrafish, despite nematodes having HSP homologs, they did not exhibit the same high expression of HSP genes as internal aging signals. The plausible explanation is that even if some aging processes follow the CARS model, the input signals it integrates likely arise from responses to different types of cellular damage, which vary depending on the cell’s function and external environment. Consequently, even within the CARS model, aging signals and the nuclear genetic Aging Program would have to adapt passively to cellular damage and error accumulation, making a fully programmed aging process unlikely.

Another reason for the illusion that aging might be a programmed process is the presence of evolutionarily conserved aging-related genes, such as those in the HSP family^93^. These genes are found across diverse species and are often implicated in stress responses and cellular maintenance^94^. However, our gene co-expression analysis and GRN analysis revealed significant differences in the transcriptional regulation of these conserved genes across species. While individual HSP genes may perform similar biological functions, how they are organized into functional modules and the timing of their expression differ greatly between species. These findings highlight that while some genes may be conserved at the sequence level, their regulatory mechanisms and roles in aging can vary dramatically. This discrepancy (conserved genes exhibiting divergent regulation) could arise from biological processes like oxidative damage, protein misfolding, DNA damage, and epigenetic changes. These processes are ancient and ubiquitous, having evolved long before the advent of complex multicellular organisms. Genes involved in countering these stresses have likely been conserved because they play fundamental roles in cellular maintenance and survival, particularly during early developmental stages.

As species diverged and adapted to different ecological niches and survival pressures, the deployment of these conserved genes in response to aging-related stresses also evolved. This indicates that aging is not controlled by a single, conserved program but rather by species-specific strategies that utilize conserved genes in different organizational frameworks. These differences in gene organization further suggest that even if an ancient programmed death mechanism did exist in a common ancestor, it would have been easily lost during evolution as aging-related genes were deployed differently across species. Given this diversity, it is possible that some organisms evolved specialized programmed aging mechanisms in response to unique environmental pressures. For instance, semelparity, a reproductive strategy seen in octopuses, Pacific salmon, and Antechinus, represents such an adaptation, but these are individual cases and do not represent a universal or conserved mechanism of programmed aging.

Finally, our results indicate the inherent diversity of aging mechanisms, which holds important implications for aging research. Since much of aging research aims to understand the underlying causes of human aging, it is crucial to prioritize model organisms more closely related to humans when studying complex aging processes. This is especially relevant when attempting to elucidate the transcriptional regulatory networks and molecular pathways involved in human aging, as these are more likely to reflect the specific aging mechanisms relevant to our species. In conclusion, our findings suggest that aging is not controlled by a conserved, ancient program, but rather shaped by species-specific adaptations. This insight has significant implications for our understanding of aging mechanisms and the development of interventions to modulate aging and extend health span across diverse organisms.

## Methods

### Data Acquisition and Preprocessing

We retrieved publicly available single-cell transcriptomic datasets of aging from zebrafish, fruit fly, and nematode, each dataset spanning six distinct time points representing various stages of adulthood. The single-cell transcriptomic expression matrix file for zebrafish was downloaded from https://figshare.com/s/1ab3c6d7648d12247eb2, where we obtained the ZDCL.rdata file. We also downloaded the cell annotation information from https://bis.zju.edu.cn/ZCL/. For fruit fly, we downloaded the single-cell transcriptomic expression matrices of both the head and body, along with the cell annotation information, from the GEO database (accession number GSE218661).

To ensure consistency across datasets, we applied log normalization to stabilize the variance in gene expression values. Dimensionality reduction was performed using UMAP and PCA, and the cell type annotations were mapped to the gene expression matrices. All processed datasets were saved in rds and h5ad file formats using Seurat and Scanpy for efficient downstream analysis. The single-cell transcriptomic data for nematodes was downloaded from “The Complete Cell Atlas of Nematode Aging” at https://c.elegans.aging.atlas.research.calicolabs.com/data. We downloaded the h5ad file and converted it into the rds format after standardization.

### Cell Type Selection and Single-cell Co-expression Module Identification

Cell type distributions across all datasets were analyzed, and representative cell types with adequate numbers were selected for downstream analysis. Biologically significant cell types were prioritized, including epithelial cells, germ cells, glial cells, neurons, muscle cells, and seam cells. These cell types were selected from each organism to ensure their biological relevance to aging studies.

We used hdWGCNA (high-dimensional Weighted Gene Co-expression Network Analysis) to perform WGCNA on the selected cell types across different life stages for each species. To construct the network, we chose the soft-thresholding power that ensured a scale-free topology, setting a minimum R² ≥ 0.8. The maximum module size was set to 30,000 genes and the minimum module size to 30 genes to create robust co-expression networks. For each gene within the network, KME (module eigengene connectivity) values were calculated. Genes with KME > 0.6 were classified as hub genes, while genes with KME > 0.4 were included in the overall gene set of the module.

### Comparative Analysis Across Tissues

To compare co-expression modules across tissues and species (zebrafish, fruit fly, and nematode), we employed various tools and software to ensure accuracy and consistency in our analyses. For Venn diagram generation, we used the *VennDiagram* package in R, leveraging the draw.triple.venn() function to compare the top 30 genes from each module across tissues. A kME threshold greater than 0.4 was used to select significant genes, ensuring that only key contributors to each module were included in the comparisons. Additionally, we utilized *UpSetR* to visualize overlaps across multiple tissues, allowing for multi-tissue comparisons of co-expression modules across the species. For enrichment analysis, we utilized *FishEnrichr*, *FlyEnrichr*, and *NematodeEnrichr* tools, performing KEGG 2019 and GO Biological Process 2018 analyses. Genes from each co-expression module (top 100 by kME) were submitted, and the Z-scores and adjusted p-values were calculated to identify pathways enriched in aging, stress responses, and cellular maintenance. The results were visualized using *ggplot2*, with bar plots highlighting the most enriched pathways per module. To complement this, we performed differential expression analysis using the *DESeq2* package in R, focusing on heat shock protein (HSP) genes across nematode tissues and life stages (days 1, 3, 5, 8, 11, and 15). Raw counts were normalized, and fold changes were assessed, with significant differential expression defined as an adjusted p-value < 0.05 and a fold change > 2.

### Gene Enrichment Analysis and Differential Expression Analysis

For enrichment analysis, we utilized *FishEnrichr*, *FlyEnrichr*, and *NematodeEnrichr* tools, performing KEGG 2019 and GO Biological Process 2018 analyses. Genes from each co-expression module (top 100 by kME) were submitted, and the Z-scores and adjusted p-values were calculated to identify pathways enriched in aging, stress responses, and cellular maintenance. The results were visualized using *ggplot2*, with bar plots highlighting the most enriched pathways per module^95, 96^.To complement this, we performed differential expression analysis using the *DESeq2* package in R, focusing on heat shock protein (HSP) genes across nematode tissues and life stages (days 1, 3, 5, 8, 11, and 15). Raw counts were normalized, and fold changes were assessed, with significant differential expression defined as an adjusted p-value < 0.05 and a fold change > 2.

### Gene Regulatory Network Construction

We used CellOracle to construct Gene Regulatory Networks (GRNs) for each species^97^. The processed H5AD files (containing normalized and dimensionally reduced data via UMAP and PCA) were used to infer GRNs using *GRNBoost2* within *CellOracle*, with transcription factors scoring a Gini importance greater than 0.1 retained. Cross-species comparisons of the GRNs were made by identifying homologous transcription factor-target pairs using the *Alliance of Genome Resources* database. For zebrafish, we focused on the RPS/RPL and HSP-UPS modules, extracting the top 10 genes by kME from each module to form local subnetworks. These were visualized using *Cytoscape*, where transcription factors were displayed as green squares and effector proteins as blue circles. In addition to visualization, we performed connectivity analysis using *igraph* and *networkx*, calculating the node degree, betweenness centrality, and clustering coefficients to assess the regulatory complexity of the subnetworks^98^. Cross-species comparisons of the RPS/RPL and HSP-UPS GRNs enabled us to assess the conservation of transcriptional regulatory mechanisms in aging processes.

To further compare the conservation of these networks, we mapped orthologous genes across zebrafish, fruit fly, and nematode, identifying homologous transcriptional regulators for both the HSP-UPS and RPS/RPL modules. For the zebrafish GRNs, six transcription factors (foxa3, foxp, cdc5l, lhx5, hnf4a, and erg) co-regulated 13 ribosomal protein-coding genes, with similar co-regulation patterns observed in fruit fly (fkh, fd68A, fd19B, bin) and nematode (sox-2, nhr-35, lin-11, fkh-7, ast-1, pha-4). These comparisons enabled us to assess the conservation of gene regulatory mechanisms in aging across different species.

