## Supplementary Figs. 1-3 and Tables 1 for "Cross-Species Analysis Reveals No Universal Programmed Aging Mechanism: Insights from Single-Cell Transcriptomics in Zebrafish, Fruit Fly, and Nematode"

**Supplementary Information**

**Supplementary Figures 1-3**

**Tables S1**

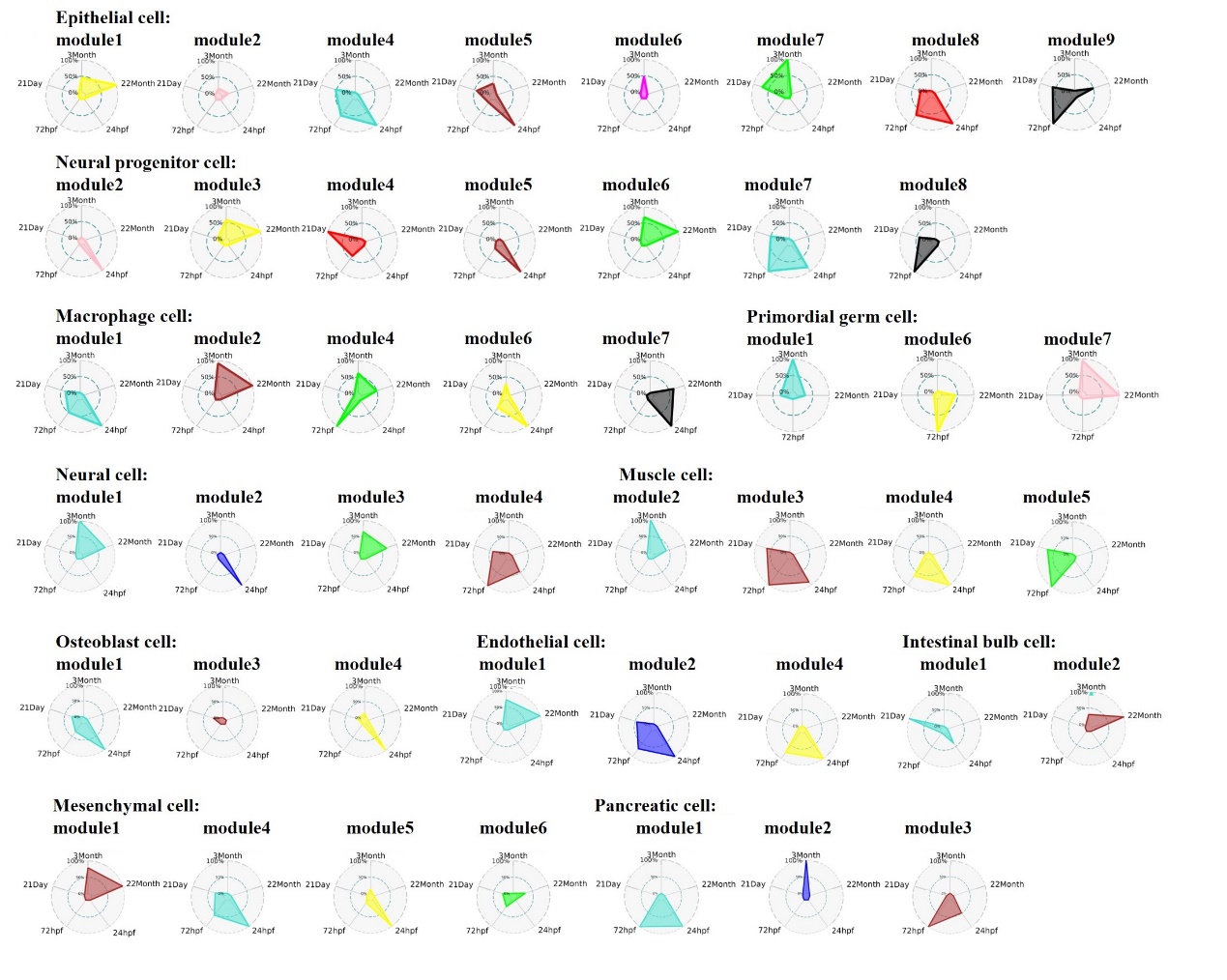

**Supplementary Figure 1 The expression changes of all co-expression modules across 11 tissues in zebrafish with aging.**

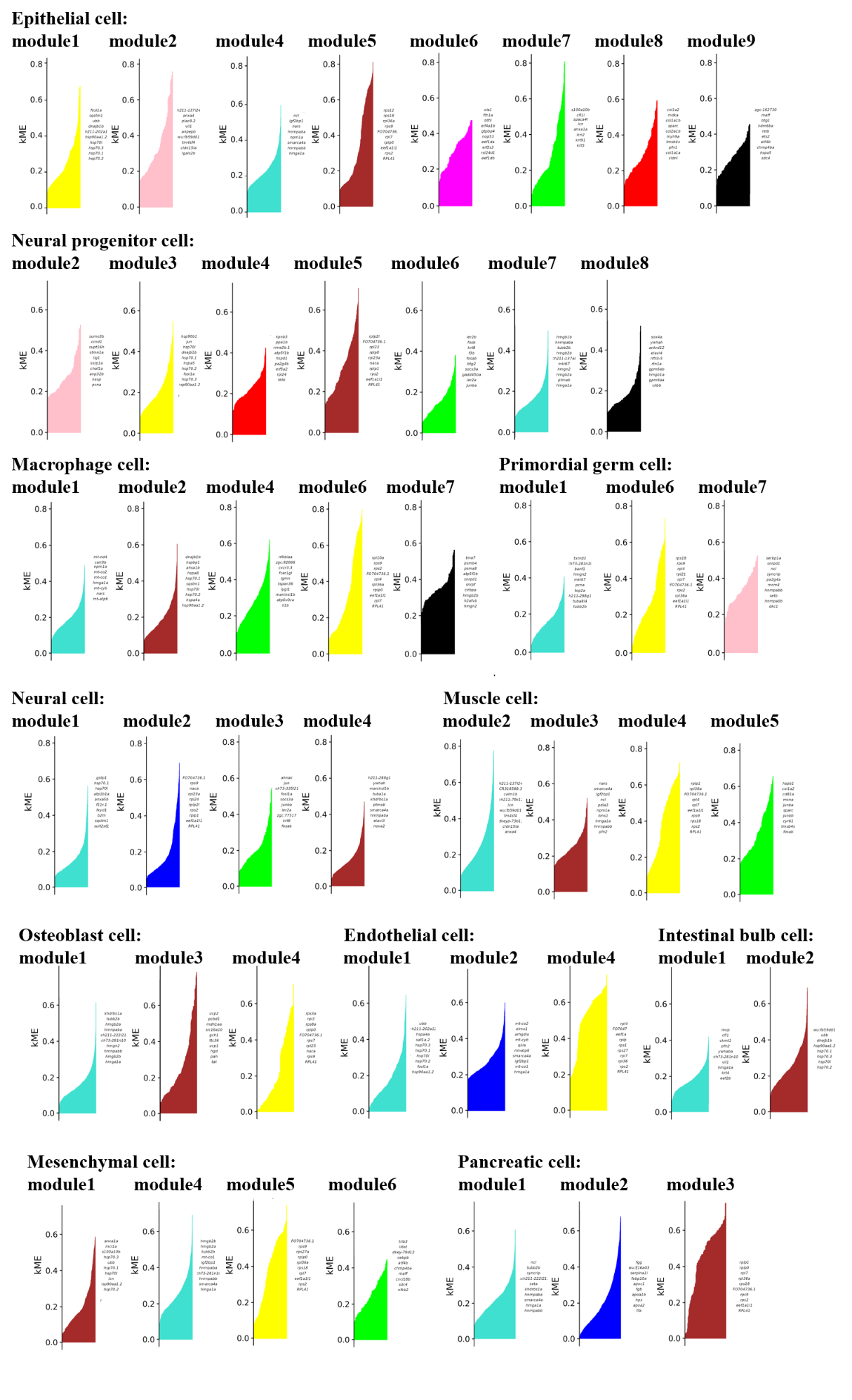

**Supplementary Figure 2 The kME distribution across all co-expression modules in the 11 zebrafish tissues, along with the top 10 genes with the highest kME values within each module.**

**Table S1. The homology and functional prediction of genes involved in the construction of GRNs in zebrafish, fruit fly, and nematode.**

| **Zebrafish** | **Drosophila** | **C. elegans** | **Gene Description** |
| --- | --- | --- | --- |
| fosl1a | kay | / | Transcription factor involved in RNA polymerase II regulation; acts in transcriptional regulation, expressed in various tissues. |
| ets2 | pnt | / | Transcription factor, involved in cell differentiation and transcription regulation; expressed in various developmental structures. |
| bhlhe40 | / | / | Regulates circadian gene expression and transcription; involved in rhythmic behavior and expressed in various systems. |
| jdp2b | Atf3 | / | Transcription factor involved in transcriptional regulation; localized to the nucleus and active in several processes. |
| junba | jra | 1-Jun | Transcription factor in RNA polymerase II regulation; plays roles in heart and lymph vessel development, among other processes. |
| junbb | jra | 1-Jun | Similar to junba, involved in transcriptional regulation and various developmental processes. |
| jund | jra | 1-Jun | Transcription factor in cell cycle and cell proliferation regulation, expressed in telencephalon. |
| cebpb | slbo | cebp-1, zip-4 | Transcription factor involved in bacterial response and gene regulation; active in adipose tissue and macrophages. |
| nr1d2b | Eip75B | nhr-85, sex-1 | Transcription factor in hormone signaling and cell differentiation; expressed in nervous system and optic vesicle. |
| fosb | / | / | Transcription factor involved in gene regulation, expressed in brain and epidermis. |
| nfe2l2a | cnc | skn-1 | Transcription factor, involved in cellular responses and DNA binding; expressed in the digestive system and other organs. |
| nr4a3 | Hr38 | nhr-6 | Transcription factor involved in glucocorticoid receptor binding and neutrophil regulation. |
| bach1b | CG15725 | / | Transcription factor involved in gene regulation; expressed in blood, pancreas, and nervous tissues. |
| maff | maf-S | maf-1 | Transcription factor involved in DNA binding and transcription regulation; forms part of regulatory complexes. |
| slc25a25a | SCaMC | F17E5.2 | Involved in ATP/ADP transport across membranes, expressed in brain, ear, and somites. |
| ubb | Ubi-p63E | ubq-1 | Ubiquitin pathway gene, involved in protein degradation processes. |
| sat1a.2 | CG4210 | D2023.4 | Involved in spermidine acetylation, expressed in digestive and other systems. |
| hsp70.3 | Hsc70-1, Hsc70-4 | F26D10.24 | Heat shock protein involved in protein folding and stress responses. |
| hsp70.2 | Hsc70-1, Hsc70-4 | F26D10.24 | Heat shock protein involved in protein folding and stress responses. |
| hsp70.1 | Hsc70-1, Hsc70-4 | F26D10.24 | Heat shock protein involved in protein folding and stress responses. |
| hsp90aa1.2 | Hsp83 | hsp-90 | Heat shock protein involved in protein stabilization and folding. |
| hsp70l | Hsc70-1, Hsc70-4 | F26D10.24 | Heat shock protein involved in protein folding and stress responses. |
| Sox2 | Sox100B, bin | sox-2 | Chromatin and sequence-specific DNA-binding transcription factor; involved in fin regeneration, nervous system development, and otic placode development. |
| cdc5l | / | cdc-5L | Transcription factor involved in mRNA splicing and RNA polymerase II regulation; associated with osteosarcoma. |
| hnf4a | / | nhr-35 | Transcription factor involved in cell differentiation and transcription regulation; related to type 2 diabetes mellitus. |
| lhx5 | / | lin-11 | Transcription factor involved in axonal fasciculation and positive regulation of vulval development. |
| foxp4 | bin | fkh-7 | Transcription factor involved in transcriptional regulation and development processes. |
| erg | Ets21C | ast-1 | Transcription factor involved in neuron differentiation and transcription regulation. |
| foxa3 | fkh, fd68A, fd19B | pha-4 | DNA-binding transcription factor; plays a role in lifespan determination, sodium arsenite response, and macromolecule biosynthesis. |
| rpl3 | RpL3 | rpl-3 | A structural constituent of ribosomes; involved in cytoplasmic translation. |
| rps3a | RpS3A | rps-1 (rps3A) | Structural constituent of ribosome, involved in cytoplasmic translation. |
| rpl18 | RpL18 | rpl20 (rpl18) | Ribosomal protein, part of the ribosomal large subunit involved in translation. |
| rpl13a | RpL13A | rpl-16 (rpl-13A) | Ribosomal protein involved in negative regulation of translation. |
| rpl10 | RpL10 | rpl-10 (rpl-10L) | Ribosomal protein involved in brain development and translation regulation. |
| rpl10a | RpL10Ab | rpl-1 (rpl-10a) | Ribosomal protein involved in T cell differentiation and chordate embryonic development. |
| rpl13 | RpL13 | rpl-13 | Ribosomal protein involved in translation. |
| rps7 | RpS7 | rps-7 | Structural component of ribosomes, involved in rRNA processing and ribosomal subunit biogenesis. |
| rps9 | RpS9 | rps-9 | Ribosomal protein involved in the structural formation of ribosomes. |
| rps23 | RpS23 | rps-23 | A key ribosomal protein involved in translation. |
| rpl9 | RpL9 | rpl-9 | Ribosomal protein involved in protein synthesis and translational regulation. |
| rpl31 | RpL31 | rpl-31 | Ribosomal protein involved in protein synthesis. |
| xbp1 | Xbp1 | xbp-1 | Transcription factor involved in unfolded protein response and stress management. |
